# Metabolic Drivers of Disease Activity and Complications in Crohn’s Disease: A Retrospective Cross-Sectional Study

**DOI:** 10.64898/2026.04.01.715942

**Authors:** Yu-Xia Pan, Shan-Shan Huang, Shan-Yu Qin, Zhu-Dong Liu, Yan-Hua Liang, Hai-Xing Jiang

**Affiliations:** Guangxi Medical University, No. 22 Shuangyong Road, Nanning 530021, Guangxi, China; The First Affiliated Hospital of Guangxi Medical University, No. 6 Shuangyong Road, Nanning 530021, Guangxi, China

**Keywords:** Crohn’s disease, Metabolic syndrome, Crohn’s Disease Activity Index, Intestinal complications, Linear models, Logistic models

## Abstract

**Background:** This study aims to examine the independent relationships between individual components of metabolic syndrome (MetS) and two key clinical outcomes in patients with Crohn’s disease (CD): disease activity, as quantified by the Crohn’s Disease Activity Index (CDAI), and the occurrence of complications.

**Methods:** This retrospective cross-sectional study included 376 adults with newly diagnosed Crohn’s disease. Multiple linear regression was used to examine associations between metabolic parameters and CDAI scores, while multivariate logistic regression assessed links to complications. Analyses were also based on clinical CDAI cut-offs. Predictive nomograms were developed and internally validated via bootstrap resampling.

**Results:** Multiple linear regression indicated that higher CDAI scores were independently associated with lower BMI (B = −5.866, P < 0.001), lower HDL-C levels (B = −81.770, P < 0.001), higher triglycerides (B = 15.618, P = 0.001), and lower ESR (B = −0.375, P = 0.03). Multivariate logistic regression established low HDL-C (OR = 0.042, P < 0.001), low BMI (OR = 0.915, P = 0.034), and high triglycerides (OR = 1.792, P = 0.007) as significant independent risk factors for complications. The developed nomograms demonstrated strong predictive performance, with an adjusted R^2^ of 0.207 for the CDAI model and an AUC of 0.765 for the complication model. For both predictive tasks, the model incorporating separate TG and HDL-C measurements significantly outperformed the TG/HDL-C ratio model.

**Conclusion:** Metabolic disturbances demonstrate a significant association with increased disease severity and a higher risk of complication development in Crohn’s disease.

**Core tip:**

1. Dual-outcome study reveals HDL-C and TG differentially link to CD inflammation and complications, pointing to distinct mechanisms.
2. Low HDL-C is the strongest independent predictor for CD complications, underscoring its protective role beyond cholesterol transport.
3. Individual TG and HDL-C metrics outperform their ratio in prediction, challenging its use and suggesting independent pathways in CD.
4. Low BMI independently associates with both adverse outcomes, refining the “obesity paradox” and highlighting malnutrition’s key role.
5. A practical, validated nomogram (AUC=0.765) integrates HDL-C, TG, and BMI to stratify complication risk, aiding clinical decision-making.

## 1. Introduction

Crohn’s disease (CD) is a chronic condition driven by immune dysfunction, manifesting as transmural inflammation of the intestinal tract. Its etiology is multifaceted, and patient outcomes often vary considerably^1^. Persistent transmural injury results in irreversible intestinal fibrosis. Over time, the combined effects of chronic inflammation and fibrotic remodeling contribute to progressive bowel wall thickening, the development of strictures, and complications such as obstruction or penetration^2^. Clinically, this heterogeneity is observed across two distinct sides: short-term disease activity, typically measured by tools such as the Crohn’s Disease Activity Index (CDAI), and long-term structural sequelae, including strictures, fistulas, and abscesses. Despite the therapeutic advances brought by biologic agents, significant variability in patient outcomes persists. A subset of individuals continues to develop refractory inflammatory responses, fibrotic strictures, and penetrating disease complications^3^. We urgently need to find a new biomarker that can predict both Disease activity and risk of complications.

Over recent decades, the prevalence of both metabolic syndrome (MetS) and inflammatory bowel disease (IBD) has increased in parallel. This trend is especially evident in populations characterized by lifestyles associated with Westernization^4^. A Western-style dietary pattern can promote intestinal inflammation by activating innate immune receptors and disrupting the gut microbiota^5^. The state of obesity represents an established risk factor for inflammatory bowel disease (IBD). This association is mediated in part by underlying metabolic dysfunction, where chronic inflammatory processes serve as a key mechanistic pathway^6^. The conclusions of existing studies on how obesity affects the natural course of Crohn’s disease (CD) are inconsistent, reflecting a complex association between the two that has not been fully elucidated^7^. No studies have really elucidated the underlying pathophysiological mechanisms..

Circulating metabolic factors, especially high-density lipoprotein (HDL), are regarded as passive lipid transporters and are indispensable dynamic platforms in the immune regulatory signaling network^8^. Under normal physiological conditions, high-density lipoprotein (HDL) displays robust anti-inflammatory and antioxidant capabilities. It functions by suppressing the production of pro-inflammatory mediators from monocytes and macrophages. HDL modulates immune responses through the depletion of cholesterol within lipid rafts of antigen-presenting cells. This activity leads to the downregulation of major histocompatibility complex (MHC) class II molecules, thereby restraining excessive activation of T-lymphocytes^8^. During states of chronic inflammation, the composition and functionality of HDL are subject to substantial remodeling. Specific alterations include the incorporation of acute-phase proteins such as serum amyloid A1 and a marked reduction in its intrinsic antioxidant enzymatic activity. As a result of these changes, HDL transitions into a dysfunctional state and may even acquire distinct pro-inflammatory properties^8^. In CD, systemic metabolic disturbances may disrupt intestinal immune homeostasis by inducing dysfunction of immunometabolic products like HDL, thereby exacerbating inflammation and promoting complications. But the direct clinical evidence supporting this hypothesis is currently lacking. Therefore, this study aims to focus on circulating immunomodulatory metabolic components, integrating clinical cohort data to evaluate associations between metabolic disturbances and core CD outcomes and to construct risk prediction models. This may provide novel prognostic biomarkers and empirical evidence for understanding metabolic-immune-tissue repair crosstalk in intestinal inflammation, suggesting potential therapeutic targets.

## 2. Subjects and Methods

### 2.1 Study Design

This study used a retrospective cross-sectional study and was reported according to the STROBE Statement for observational studies.

### 2.2 Study Population

We included 376 patients first diagnosed and treated for CD at the First Affiliated Hospital of Guangxi Medical University between April 1, 2014, and June 30, 2025. Diagnosis complied with the “Chinese Guidelines for the Diagnosis and Treatment of Crohn’s Disease (2023, Guangzhou)” ^9^. Inclusion criteria: age ≥ 18 years; first diagnosis of CD at our hospital; availability of complete baseline MetS indicators and clinical data. Exclusion criteria: severely incomplete data; concomitant severe comorbidities(malignancy, liver cirrhosis, end-stage renal disease); pregnancy or lactation. The study was approved by the Medical Ethics Committee of First Affiliated Hospital of Guangxi Medical University (Approval No.:2026-E0012).

### 2.3 Outcome Definitions

This study used a dual-outcome analysis. The primary outcome variable 1 categorized patients based on CDAI score: clinical remission (CDAI < 150), mild activity (CDAI 150-220), and moderate-to-severe activity (CDAI > 220) ^10,11^. The primary outcome variable 2 categorized patients based on the presence of complications. Complications, defined per guidelines^9^, included fistula, abscess, stricture/obstruction, perianal disease, hemorrhage, perforation, and malignancy, confirmed by computed tomography enterography (CTE), magnetic resonance enterography (MRE), or endoscopy.

### 2.4 Data Collection and Variable Definitions

Data were collected from the hospital information system. Baseline characteristics: age, sex, height, weight, smoking history, time to diagnosis (symptom onset to diagnosis), disease location (Montreal classification) ^9^, disease behavior (B1/B2/B3/B2+B3) ^9^, perianal disease, surgical history, current medications (5-ASA, corticosteroids, immunomodulators, biologics). Metabolic indicators: BMI, blood pressure(BP), fasting plasma glucose (FPG)/glycated hemoglobin (HbA1c), triglycerides (TG), total cholesterol (TC), high-density lipoprotein cholesterol (HDL-C), low-density lipoprotein cholesterol (LDL-C), uric acid (UA). Calculated indicator: TG/HDL-C ratio. Other indicators: erythrocyte sedimentation rate (ESR), thyroid function, and high-sensitivity C-reactive protein (hs-CRP). Waist circumference data were unavailable retrospectively.

### 2.5 Diagnostic Definitions for Metabolic Indicators

Definitions followed the Chinese Diabetes Society guidelines ^12^: Hyperglycemia: FPG ≥ 6.1 mmol/L or 2-hour postprandial glucose ≥ 7.8 mmol/L, or diagnosed diabetes. Hypertension: BP ≥ 130/85 mmHg or diagnosed hypertension.

### 2.6 Statistical Analysis

Analyses were performed using SPSS 25.0 and R 4.5.1. Continuous variables were assessed for normality; skewed variables are reported as median (IQR) and compared using Mann-Whitney U or Kruskal-Wallis H tests. Categorical variables are reported as frequency (%) and compared using chi-square or Fisher’s exact tests. Minimal missing data (e.g., disease duration in 15 cases, 4.0%) were handled via median imputation. All analyses were performed on the imputed dataset.

Main Analysis 1 (CDAI score): Hierarchical multiple linear regression. Model 1 adjusted for clinical factors (time to diagnosis, disease behavior, location, and current treatment). Model 2 added metabolic indicators (BMI, HDL-C, UA, etc.) to Model 1; ΔR^2^ reported.

Main Analysis 2 (Complications): Hierarchical multivariate logistic regression. Model 1 adjusted for clinical factors (disease behavior, time to diagnosis, treatment). Model 2 added metabolic indicators (BMI, HDL-C, etc.) to Model 1; the change in model χ ^2^ was reported.

Model Development and Validation: Nomograms were based on the final models. Internal validation was performed via bootstrap resampling (1000 repetitions). Performance was assessed by discrimination (AUC/adjusted R²), calibration (calibration curve, Hosmer-Lemeshow test), and clinical utility (Decision Curve Analysis, DCA).

Sensitivity Analysis: Compared performance of models using the TG/HDL-C ratio vs. separate TG and HDL-C indicators. A P-value < 0.05 was considered statistically significant.

## 3. Results

### 3.1 Baseline Patient Characteristics

Among 376 CD patients, 255 (67.8%) had complications. The median CDAI score was 257 (IQR: 220-297.48). According to clinical cut-offs, 5 patients (1.3%) were in clinical remission, 98 (26.1%) had mild activity, and 273 (72.6%) had moderate-to-severe activity.

Table 1 (stratified by CDAI groups) showed significant differences (P < 0.05) for BMI, HDL-C, TC, LDL-C, smoking history, and TG/HDL-C ratio. No significant differences were observed for age, sex, time to diagnosis, disease behavior/location, surgical history, hypertension, hyperglycemia, TG, UA, ESR, hs-CRP, thyroid function, or use of biologics/corticosteroids. The proportion of patients with complications increased with higher disease activity (P < 0.001), while HDL-C (P < 0.001) and BMI (P < 0.001) decreased.

**Table 1.**
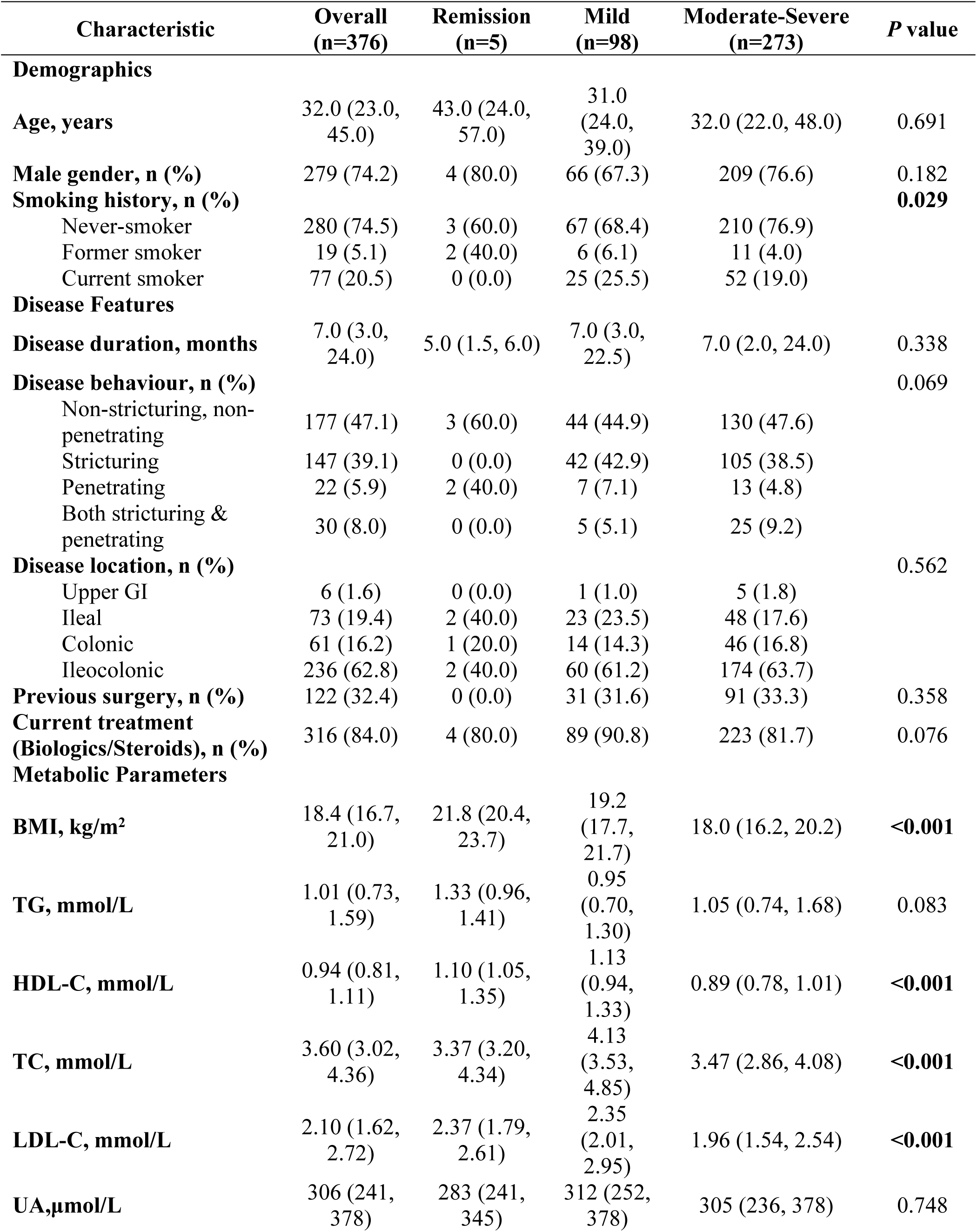
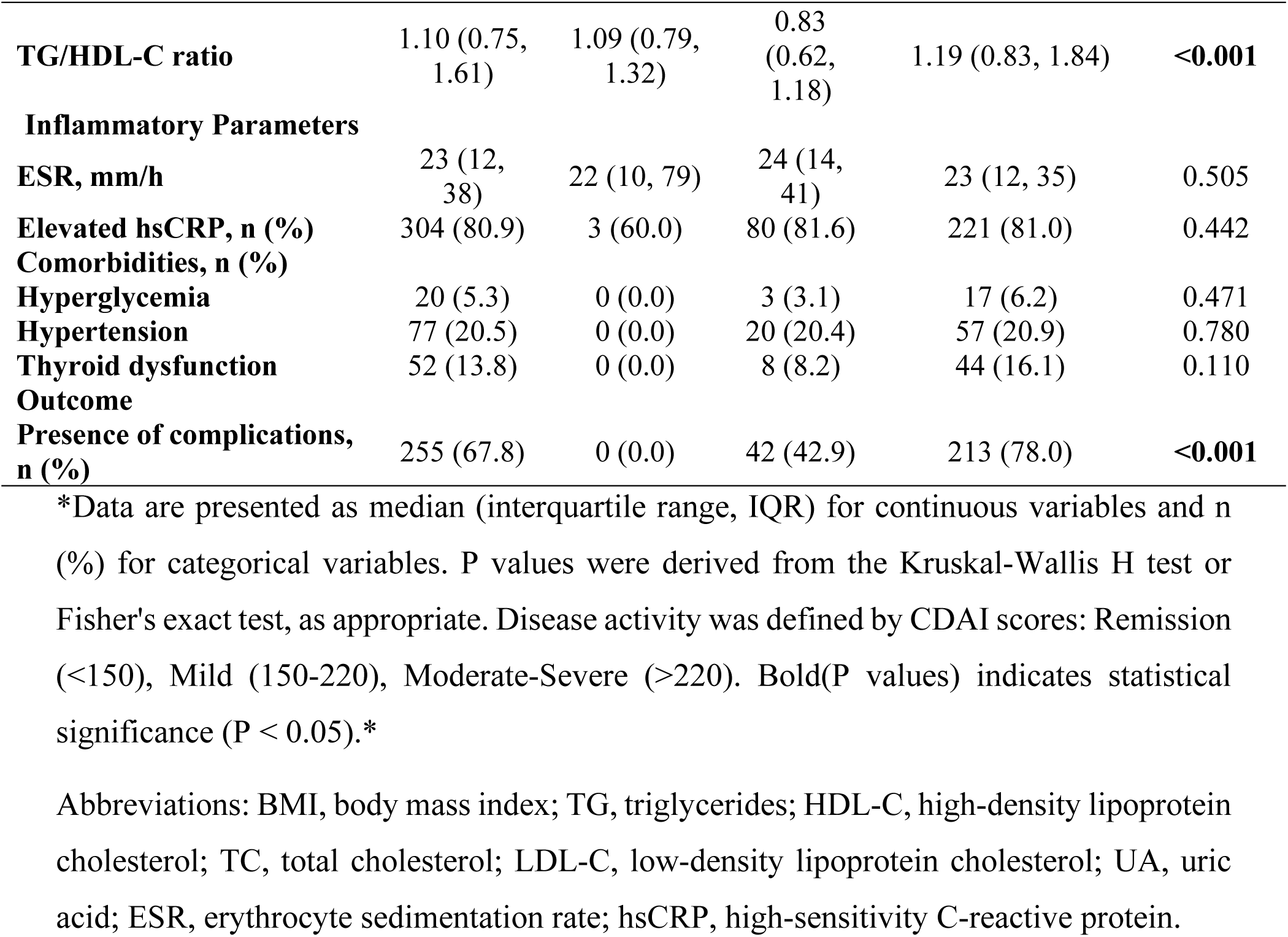
Comparison of Baseline Characteristics of All Patients Stratified by CDAI Groups.

| Characteristic | Overall<br>(n=376) | Remission<br>(n=5) | Mild<br>(n=98) | Moderate-Severe<br>(n=273) | P value |
| --- | --- | --- | --- | --- | --- |
| **Demographics** |  |  |  |  |  |
| Age, years | 32.0 (23.0, 45.0) | 43.0 (24.0, 57.0) | 31.0 (24.0, 39.0) | 32.0 (22.0, 48.0) | 0.691 |
| Male gender, n (%) | 279 (74.2) | 4 (80.0) | 66 (67.3) | 209 (76.6) | 0.182 |
| Smoking history, n (%) |  |  |  |  | **0.029** |
| Never-smoker | 280 (74.5) | 3 (60.0) | 67 (68.4) | 210 (76.9) |  |
| Former smoker | 19 (5.1) | 2 (40.0) | 6 (6.1) | 11 (4.0) |  |
| Current smoker | 77 (20.5) | 0 (0.0) | 25 (25.5) | 52 (19.0) |  |
| **Disease Features** |  |  |  |  |  |
| Disease duration, months | 7.0 (3.0, 24.0) | 5.0 (1.5, 6.0) | 7.0 (3.0, 22.5) | 7.0 (2.0, 24.0) | 0.338 |
| Disease behaviour, n (%) |  |  |  |  | 0.069 |
| Non-stricturing, non-penetrating | 177 (47.1) | 3 (60.0) | 44 (44.9) | 130 (47.6) |  |
| Stricturing | 147 (39.1) | 0 (0.0) | 42 (42.9) | 105 (38.5) |  |
| Penetrating | 22 (5.9) | 2 (40.0) | 7 (7.1) | 13 (4.8) |  |
| Both stricturing & penetrating | 30 (8.0) | 0 (0.0) | 5 (5.1) | 25 (9.2) |  |
| Disease location, n (%) |  |  |  |  | 0.562 |
| Upper GI | 6 (1.6) | 0 (0.0) | 1 (1.0) | 5 (1.8) |  |
| Ileal | 73 (19.4) | 2 (40.0) | 23 (23.5) | 48 (17.6) |  |
| Colonic | 61 (16.2) | 1 (20.0) | 14 (14.3) | 46 (16.8) |  |
| Ileocolonic | 236 (62.8) | 2 (40.0) | 60 (61.2) | 174 (63.7) |  |
| Previous surgery, n (%) | 122 (32.4) | 0 (0.0) | 31 (31.6) | 91 (33.3) | 0.358 |
| Current treatment (Biologics/Steroids), n (%) | 316 (84.0) | 4 (80.0) | 89 (90.8) | 223 (81.7) | 0.076 |
| **Metabolic Parameters** |  |  |  |  |  |
| BMI, kg/m<sup>2</sup> | 18.4 (16.7, 21.0) | 21.8 (20.4, 23.7) | 19.2 (17.7, 21.7) | 18.0 (16.2, 20.2) | **<0.001** |
| TG, mmol/L | 1.01 (0.73, 1.59) | 1.33 (0.96, 1.41) | 0.95 (0.70, 1.30) | 1.05 (0.74, 1.68) | 0.083 |
| HDL-C, mmol/L | 0.94 (0.81, 1.11) | 1.10 (1.05, 1.35) | 1.13 (0.94, 1.33) | 0.89 (0.78, 1.01) | **<0.001** |
| TC, mmol/L | 3.60 (3.02, 4.36) | 3.37 (3.20, 4.34) | 4.13 (3.53, 4.85) | 3.47 (2.86, 4.08) | **<0.001** |
| LDL-C, mmol/L | 2.10 (1.62, 2.72) | 2.37 (1.79, 2.61) | 2.35 (2.01, 2.95) | 1.96 (1.54, 2.54) | **<0.001** |
| UA, µmol/L | 306 (241, 378) | 283 (241, 345) | 312 (252, 378) | 305 (236, 378) | 0.748 |
| **TG/HDL-C ratio** | 1.10 (0.75, 1.61) | 1.09 (0.79, 1.32) | 0.83 (0.62, 1.18) | 1.19 (0.83, 1.84) | **<0.001** |
| **Inflammatory Parameters** |  |  |  |  |  |
| **ESR, mm/h** | 23 (12, 38) | 22 (10, 79) | 24 (14, 41) | 23 (12, 35) | 0.505 |
| **Elevated hsCRP, n (%)** | 304 (80.9) | 3 (60.0) | 80 (81.6) | 221 (81.0) | 0.442 |
| **Comorbidities, n (%)** |  |  |  |  |  |
| **Hyperglycemia** | 20 (5.3) | 0 (0.0) | 3 (3.1) | 17 (6.2) | 0.471 |
| **Hypertension** | 77 (20.5) | 0 (0.0) | 20 (20.4) | 57 (20.9) | 0.780 |
| **Thyroid dysfunction** | 52 (13.8) | 0 (0.0) | 8 (8.2) | 44 (16.1) | 0.110 |
| **Outcome** |  |  |  |  |  |
| **Presence of complications, n (%)** | 255 (67.8) | 0 (0.0) | 42 (42.9) | 213 (78.0) | **<0.001** |

\*Data are presented as median (interquartile range, IQR) for continuous variables and n (%) for categorical variables. P values were derived from the Kruskal-Wallis H test or Fisher's exact test, as appropriate. Disease activity was defined by CDAI scores: Remission (<150), Mild (150-220), Moderate-Severe (>220). Bold(P values) indicates statistical significance (P < 0.05).\*
Abbreviations: BMI, body mass index; TG, triglycerides; HDL-C, high-density lipoprotein cholesterol; TC, total cholesterol; LDL-C, low-density lipoprotein cholesterol; UA, uric acid; ESR, erythrocyte sedimentation rate; hsCRP, high-sensitivity C-reactive protein.

Table 2 (stratified by complication status) showed significant differences (P < 0.05) for sex, use of biologics/corticosteroids, TG, HDL-C, TC, and TG/HDL-C ratio. No significant differences were found for age, smoking history, time to diagnosis, disease behavior/location, surgical history, hypertension, hyperglycemia, BMI, LDL-C, UA, ESR, hs-CRP, or thyroid function. The complication group had a higher median CDAI (272.70 vs. 220.47, P < 0.001), lower HDL-C (P < 0.001), and a higher TG/HDL-C ratio (P < 0.001). Male sex was more frequent in the complication group (76.6% vs. 62.8%, P = 0.001).

**Table 2.** Comparison of Baseline Characteristics of All Patients Stratified by Complication Status.

| Characteristic | Overall<br>(n=376) | Without complications<br>(n=121) | With complications<br>(n=255) | P value |
| --- | --- | --- | --- | --- |
| **Demographics** |  |  |  |  |
| Age, years | 32.0 (23.0, 45.0) | 31.0 (24.0, 43.5) | 32.0 (22.0, 47.0) | 0.804 |
| Male gender, n (%) | 279 (74.2) | 76 (62.8) | 203 (79.6) | **0.001** |
| Smoking history, n (%) |  |  |  | 0.256 |
| Never-smoker | 280 (74.5) | 85 (70.2) | 195 (76.5) |  |
| Former smoker | 19 (5.1) | 9 (7.4) | 10 (3.9) |  |
| Current smoker | 77 (20.5) | 27 (22.3) | 50 (19.6) |  |
| **Disease Features** |  |  |  |  |
| Disease duration, months | 7.0 (3.0, 24.0) | 6.0 (2.0, 12.0) | 9.0 (3.0, 24.0) | 0.055 |
| Disease behaviour, n (%) |  |  |  | 0.321 |
| Non-stricturing, non-penetrating | 177 (47.1) | 54 (44.6) | 123 (48.2) |  |
| Stricturing | 147 (39.1) | 47 (38.8) | 100 (39.2) |  |
| Penetrating | 22 (5.9) | 11 (9.1) | 11 (4.3) |  |
| Both stricturing & penetrating | 30 (8.0) | 9 (7.4) | 21 (8.2) |  |
| Disease location<sup>a</sup>, n (%) |  |  |  | 0.090 |
| Upper GI | 6 (1.6) | 0 (0.0) | 6 (2.4) |  |
| Ileal | 73 (19.4) | 28 (23.1) | 45 (17.6) |  |
| Colonic | 61 (16.2) | 24 (19.8) | 37 (14.5) |  |
| Ileocolonic | 236 (62.8) | 69 (57.0) | 167 (65.5) |  |
| Previous surgery, n (%) | 122 (32.4) | 33 (27.3) | 89 (34.9) | 0.140 |
| Current treatment (Biologics/Steroids), n (%) | 316 (84.0) | 109 (90.1) | 207 (81.2) | **0.028** |
| **Metabolic Parameters** |  |  |  |  |
| BMI, kg/m<sup>2</sup> | 18.4 (16.7, 20.9) | 18.8 (17.0, 21.3) | 18.3 (16.3, 20.7) | 0.089 |
| TG, mmol/L | 1.01 (0.73, 1.59) | 0.93 (0.71, 1.30) | 1.06 (0.75, 1.71) | **0.011** |
| HDL-C, mmol/L | 0.94 (0.81, 1.11) | 1.12 (0.93, 1.30) | 0.88 (0.77, 1.00) | **<0.001** |
| TC, mmol/L | 3.60 (3.02, 4.36) | 3.76 (3.40, 4.39) | 3.52 (2.80, 4.31) | **0.003** |
| LDL-C, mmol/L | 2.10 (1.62, 2.72) | 2.14 (1.76, 2.66) | 2.06 (1.53, 2.74) | 0.126 |
| UA, µmol/L | 306 (241, 378) | 298(241, 338) | 312(240, 392) | 0.133 |
| TG/HDL-C ratio | 1.10 (0.75, 1.61) | 0.86 (0.63, 1.32) | 1.19 (0.84, 1.98) | **<0.001** |
| **Inflammatory Parameters** |  |  |  |  |
| ESR, mm/h | 23(12, 38) | 23(12, 39) | 23.0 (12, 38) | 0.514 |
| Elevated hsCRP, n (%) | 304 (80.9) | 94 (30.9) | 210 (69.1) | 0.283 |
| **Comorbidities, n (%)** |  |  |  |  |
| Hyperglycemia | 20 (5.3) | 3 (2.5) | 17 (6.7) | 0.091 |
| Hypertension | 77 (20.5) | 24 (19.8) | 53 (20.8) | 0.831 |
| Thyroid dysfunction | 52 (13.8) | 13 (10.7) | 39 (15.3) | 0.232 |
| **Disease Activity** |  |  |  |  |

| Characteristic | Overall<br>(n=376) | Without complications<br>(n=121) | With complications<br>(n=255) | *P* value |
| --- | --- | --- | --- | --- |
| **CDAI score** | 257.0<br>(220.0, 297.5) | 220.5 (187.1, 259.6) | 272.7 (237.5, 310.0) | **<0.001** |

\*Data are presented as median (interquartile range, IQR) for continuous variables and n (%) for categorical variables. P values were derived from the Mann-Whitney U test or Chi-square test, as appropriate. aDisease location's P value was derived from Fisher 's exact test. Bold(P values) indicates statistical significance (P < 0.05).\*
Abbreviations: BMI, body mass index; TG, triglycerides; HDL-C, high-density lipoprotein cholesterol; TC, total cholesterol; LDL-C, low-density lipoprotein cholesterol; UA, uric acid; ESR, erythrocyte sedimentation rate; hsCRP, high-sensitivity C-reactive protein; CDAI, Crohn's Disease Activity Index.

### 3.2 Multivariate Regression Analysis

#### 3.2.1 Factors Influencing CDAI Score

Using the CDAI score as the dependent variable, hierarchical multiple linear regression was performed. Block 1 forced entry of variables, those with a P-value of less than 0.1 in the univariate analysis (smoking history), along with clinically established important variables such as time to diagnosis, disease behavior, disease location, and current treatment. Block 2 added preset metabolic indicators (HDL-C, BMI, TG, UA, TG/HDL-C ratio, etc.) to Block 1. The incremental explanatory value of metabolic indicators was assessed via the change in R² (ΔR²). The hierarchical regression showed that (Table 3), after adjusting for clinical factors like disease duration and behavior, only four variables were independently associated with CDAI score. Lower BMI (B = −5.866, Beta = −0.258, P < 0.001), lower HDL-C (B = −81.770, Beta = −0.307, P < 0.001), higher TG (B =15.618, Beta =0.183, P =0.001), and lower ESR (B = −0.375, Beta = −0.109, P = 0.03) were independently associated with higher CDAI. Notably, traditional clinical factors lost significance after adjusting for metabolic indicators.

**Table 3:**
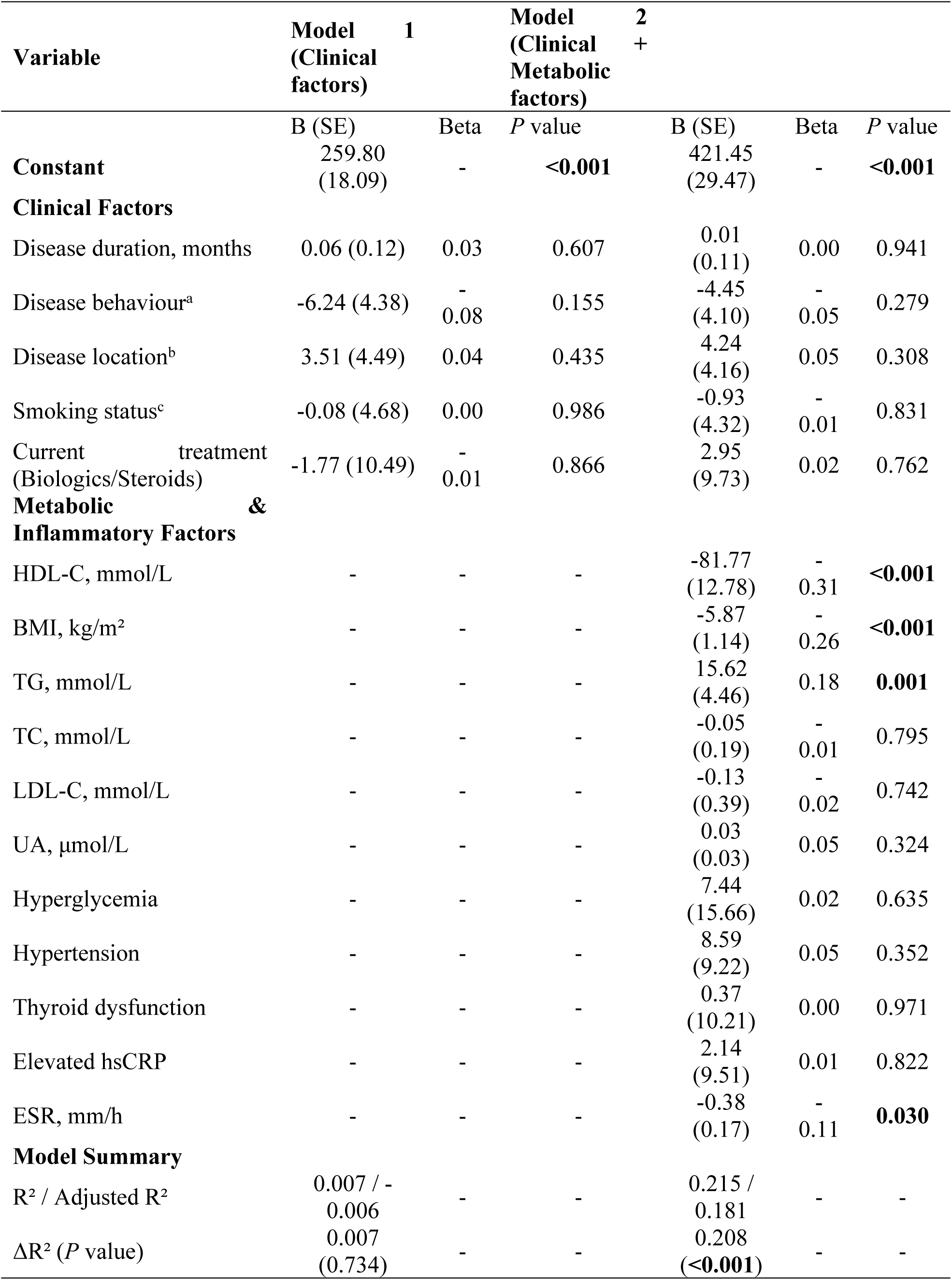
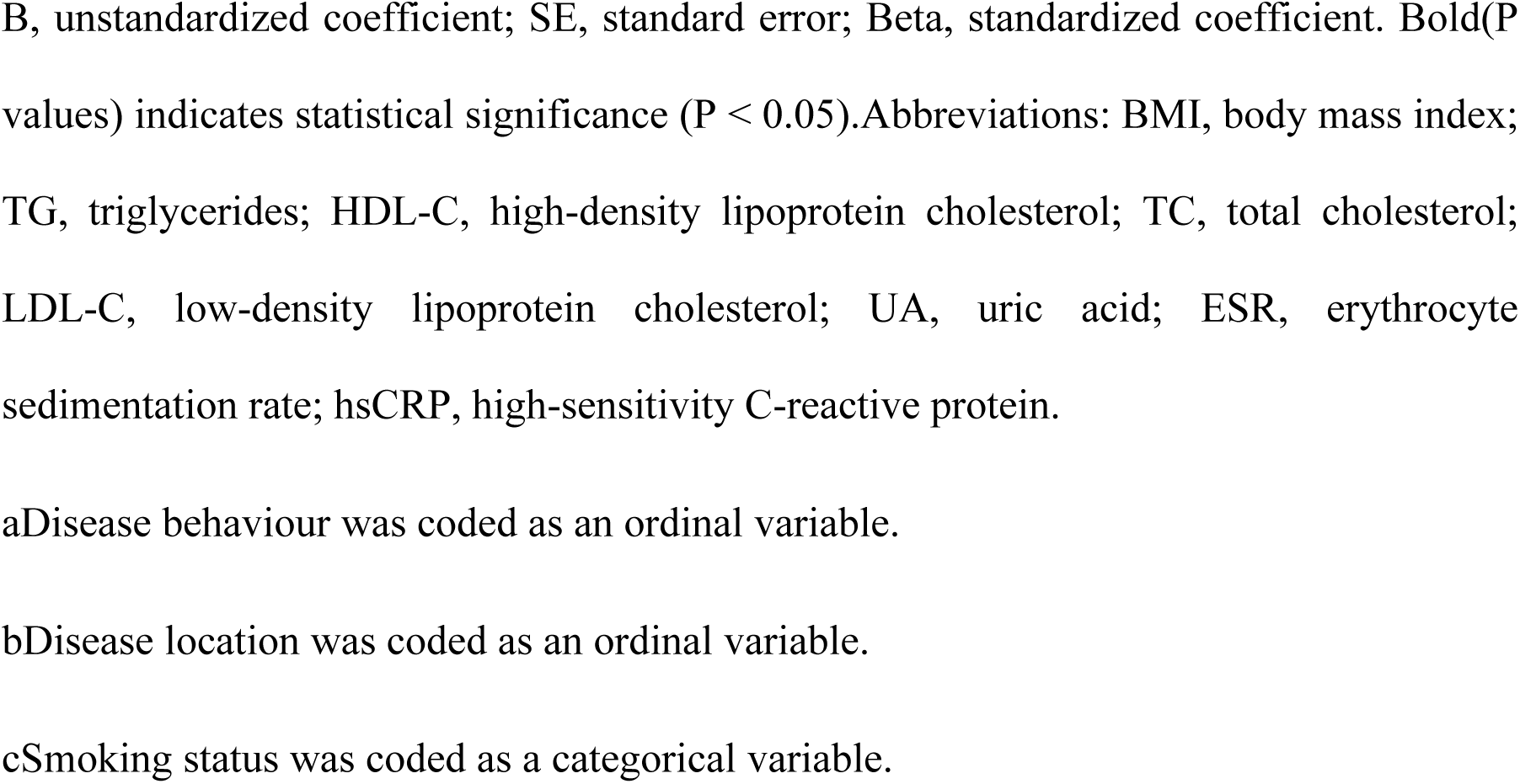
Hierarchical Linear Regression Analysis for CDAI Score.

#### 3.2.2 Independent Risk Factors for Complications

Using complication status (no=0, yes=1) as the dependent variable, hierarchical binary logistic regression was used. Due to issues of perfect separation for “disease location” identified in preliminary analysis and multicollinearity among variables, a streamlined multivariate logistic regression model was constructed. Block 1 forced entry of known important clinical variables (disease behavior, time to diagnosis, treatment) and variables with a P-value of less than 0.1 in the univariate analysis (sex). Block 2 added five metabolic/inflammatory indicators (BMI, HDL-C, TG, ESR, UA) to Block 1, focusing on changes in pseudo R², likelihood ratio tests, and the ORs with 95% CIs of metabolic indicators. Hierarchical logistic regression revealed that HDL-C (OR=0.042, 95%CI: 0.015-0.116, P<0.001) and BMI (OR=0.915, 95%CI: 0.842-0.993, P=0.034) were independent protective factors, while TG (OR=1.792, 95%CI: 1.170-2.745, P=0.007) was an independent risk factor (Table 4). Sex, time to diagnosis, disease behavior, treatment, UA, and ESR showed no independent predictive value after adjusting for metabolic factors.

**Table 4:** Multivariate Binary Logistic Regression Analysis for Complication Status.

| Variable | Model 1<br>(Clinical factors) |  | Model 2<br>(Clinical + Metabolic factors) |  |
| --- | --- | --- | --- | --- |
|  | OR (95% CI) | *P* value | OR (95% CI) | *P* value |
| **Clinical Factors** |  |  |  |  |
| Male gender | 0.48 (0.30 - 0.78) | **0.003** | 0.66 (0.37 - 1.17) | 0.15 |
| Disease behaviour | - | 0.465 | - | 0.513 |
| Disease duration, months | 1.01 (1.00 - 1.02) | 0.062 | 1.01 (1.00 - 1.02) | 0.094 |
| Current treatment<br>(Biologics/Steroids) | 0.52 (0.26 - 1.04) | 0.065 | 0.56 (0.26 - 1.21) | 0.143 |
| **Metabolic & Inflammatory Factors** |  |  |  |  |
| HDL-C, mmol/L | - | - | 0.04 (0.02 - 0.12) | **<0.001** |
| BMI, kg/m<sup>2</sup> | - | - | 0.92 (0.84 - 0.99) | **0.034** |
| TG, mmol/L | - | - | 1.79 (1.17 - 2.75) | **0.007** |
| UA, µmol/L | - | - | 1.00 (1.00 - 1.00) | 0.192 |
| ESR, mm/h | - | - | 1.00 (0.99 - 1.01) | 0.484 |
| **Model Fit Statistics** |  |  |  |  |
| -2 Log likelihood | 450.259 |  | 385.362 |  |
| Cox & Snell R<sup>2</sup> | 0.057 |  | 0.207 |  |
| Nagelkerke R<sup>2</sup> | 0.080 |  | 0.289 |  |
| Model $\chi^2$ (*P* value) | 22.166 (**0.001**) | | 87.063 (**<0.001**) | |
| $\chi^2$ change (*P* value) | - | | 64.897 (**<0.001**) | |
| Hosmer-Lemeshow test (*P* value) | 3.681 (0.885) |  | 12.761 (0.120) |  |

OR, odds ratio; CI, confidence interval. Bold(*P* values) indicates statistical significance (*P* < 0.05). Note: Disease behaviour was included in the model as a set of dummy variables; the overall *P* value is reported. Abbreviations: BMI, body mass index; TG, triglycerides; HDL-C, high-density lipoprotein cholesterol; UA, uric acid; ESR, erythrocyte sedimentation rate.

### 3.3 ROC Curves and Nomogram Models

Based on the multivariate logistic regression results, an ROC curve was constructed for complication status. The prediction model’s AUC was 0.765 (95%CI: 0.712∼0.818). Based on the Youden index, the optimal cut-off was 0.612 (sensitivity 85.9%, specificity 60.3%), with a maximum Youden index of 0.462 (Figure 1). R software validation showed consistency with SPSS results(HDL-C: SPSS=0.042 vs R=0.037; TG: SPSS=1.792 vs R=1.805; BMI: SPSS=0.915 vs R=0.922). The nomogram was constructed using R results (Figure 2). The calibration curve indicated good calibration, with a mean absolute error of 0.04. After bias correction via 1000 bootstrap resamples, the predicted probabilities remained largely consistent with observed probabilities across most ranges (Figure 3).

**Figure 1.**
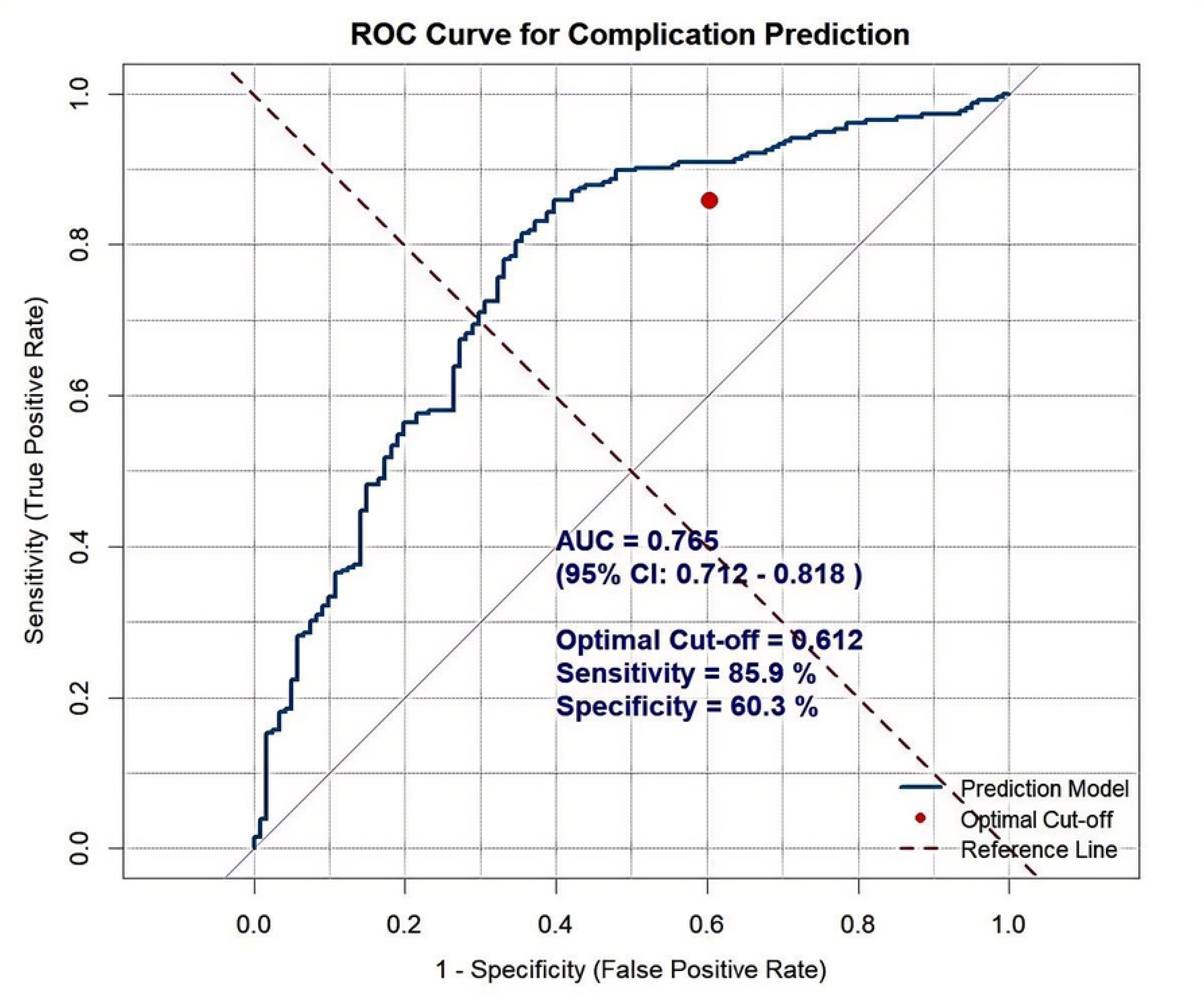
Receiver operating characteristic (ROC) curve for the complication prediction model. The area under the curve (AUC) was 0.765 (95% confidence interval [CI]: 0.712–0.818), indicating good discriminatory ability. The dotted diagonal line represents the reference line of no discrimination (AUC = 0.5).

**Figure 2.**
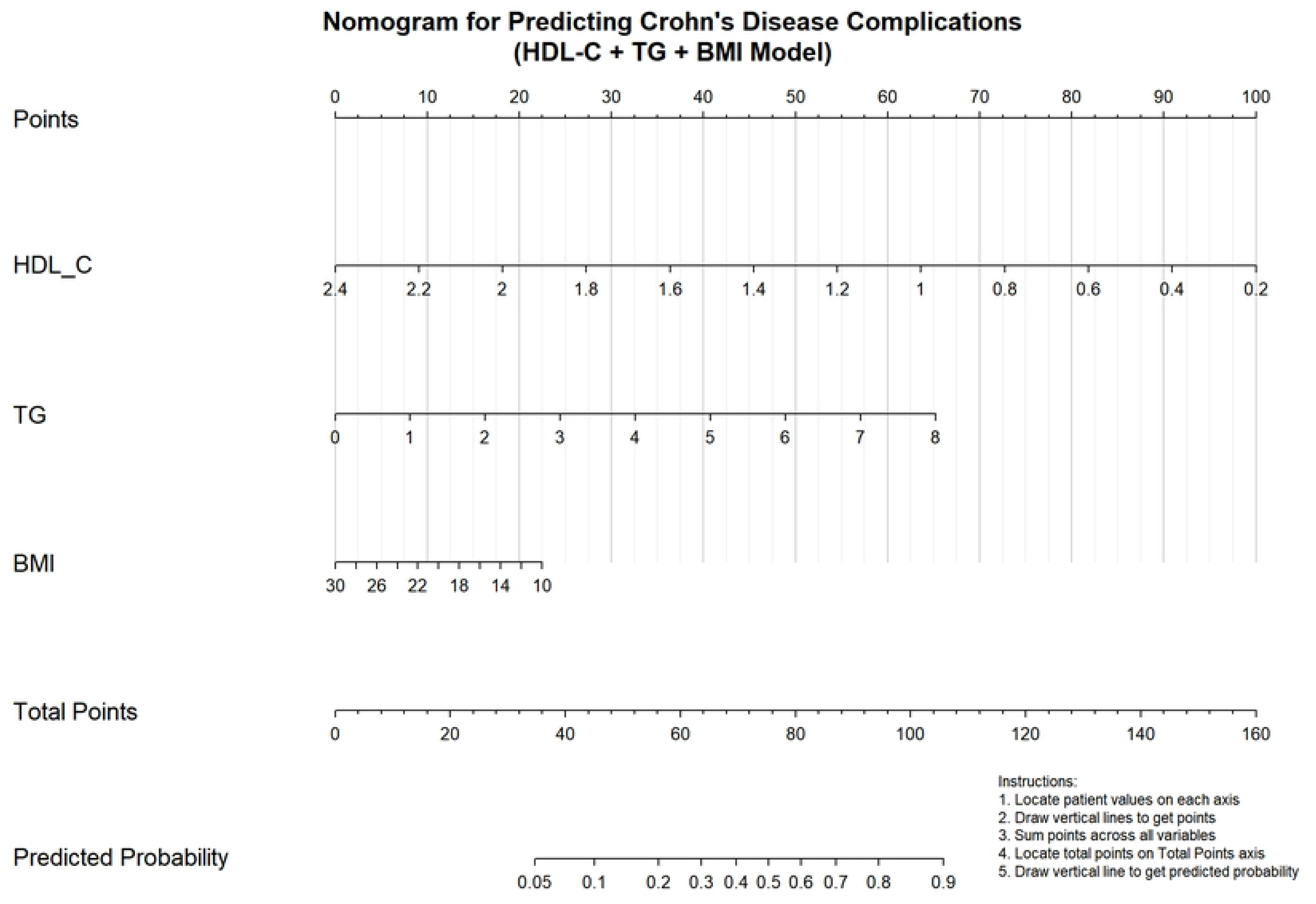
Nomogram for predicting Crohn’s disease complications. This nomogram was developed based on the final multivariate logistic regression model, incorporating high-density lipoprotein cholesterol (HDL-C), body mass index (BMI), and triglycerides (TG). To estimate the risk, locate the patient’s value for each variable on the corresponding axis, draw a line upward to the ‘Points’ line to obtain the score, sum all scores, and project the total points on the ‘Total Points’ axis down to the ‘Risk of Complication’ axis to read the predicted probability.

**Figure 3.**
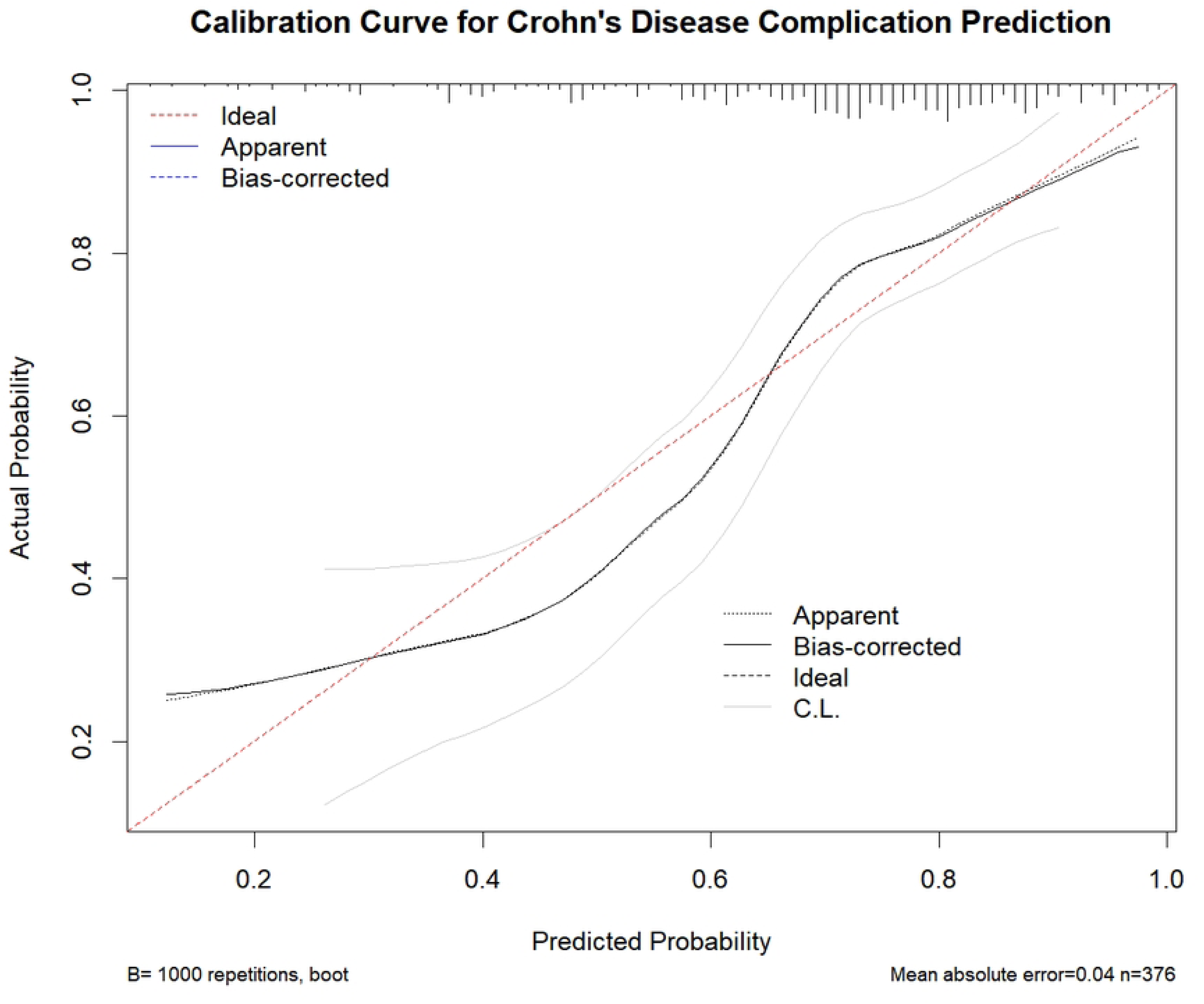
Calibration curve for Crohn’s disease complication prediction. The plot assesses the agreement between the predicted probabilities of complications (x-axis) and the observed actual frequencies (y-axis). The solid blue line represents the performance of the model after bootstrap internal validation (1000 repetitions). The dashed diagonal line represents the ideal line of perfect calibration. The histogram shows the distribution of predicted risk scores across the patient cohort.

Decision curve analysis(DCA) showed a favorable clinical net benefit across a 10%–40% threshold probability range, maximizing around 20% (Figure 4). This suggests the model’s value in identifying high-risk patients needing intervention while avoiding overtreatment of low-risk patients. We also used R software to establish a CDAI score prediction nomogram (R^2^: SPSS = 0.207 vs R = 0.204)(Supplementary Figure S1、S2), suitable for group-level risk trend assessment.

**Figure 4.**
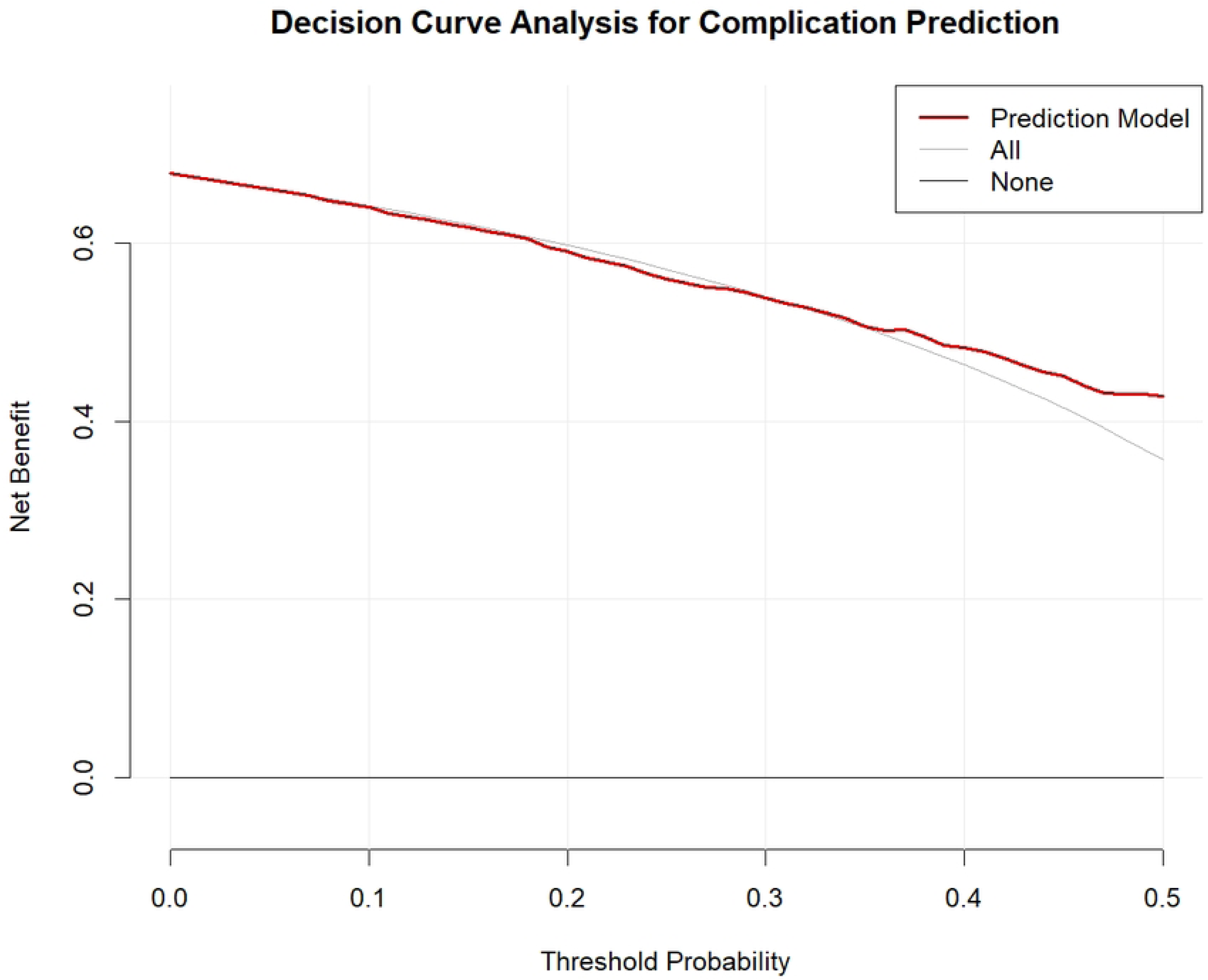
Decision Curve Analysis (DCA) for complication prediction. The net benefit of using the model (solid red line) is plotted against a range of threshold probabilities. The solid grey line and the solid black line represent the net benefit for the strategies of ‘treat all’ and ‘treat none’ patients, respectively. The model shows a positive net benefit within a clinically relevant threshold probability range of approximately 10% to 40%.

### 3.4 Sensitivity Analysis

To evaluate whether using the TG/HDL-C ratio as a composite metabolic indicator could replace separate TG and HDL-C measurements, a sensitivity analysis was performed. For the CDAI outcome, the original model’s R² (0.204) was higher than the ratio model’s R² (0.165; absolute difference ΔR² = 0.039, 95% CI: −0.0093-0.0942). For complications, the original model’s AUC (0.765) was higher than the ratio model’s (0.698; absolute difference ΔAUC = 0.0675, 95% CI: 0.0326-0.1168) (Figure 5). Comparing ROC curves stratified by complication status confirmed the superior discriminative ability of the original model (Figure 6). The original model also had lower AIC/BIC values, indicating better statistical fit and parsimony (Figure 7). Bootstrap repetition (1000 times) further confirmed the performance advantage of the original model (mean AUC difference: 0.071, 95% CI: 0.033-0.116). We found that the variation in the TG/HDL-C ratio primarily reflected TG levels (Figure 8). The original model was significantly better than the model using the TG / HDL-C ratio.

**Figure 5.**
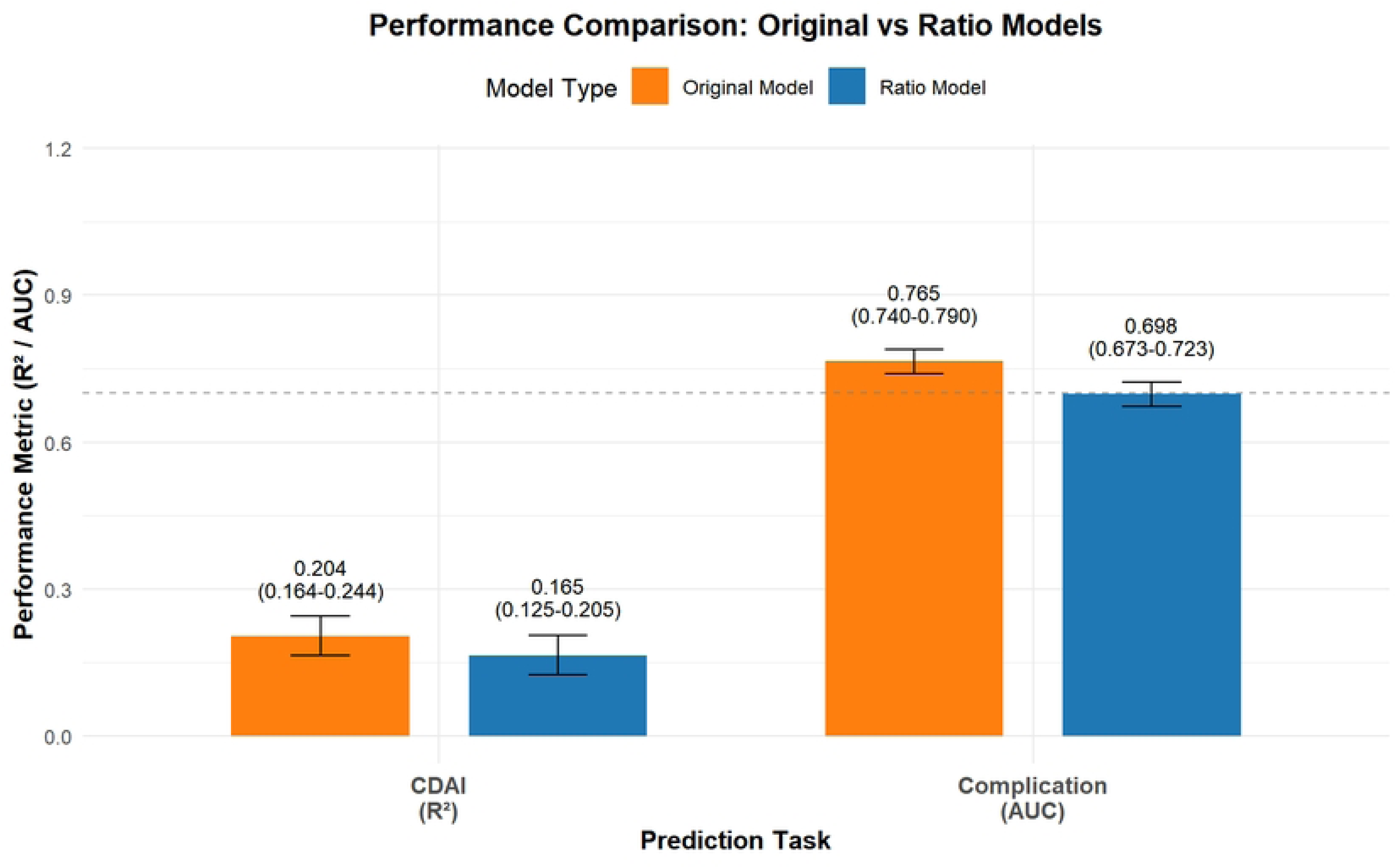
Performance comparison between the original model and the TG/HDL-C ratio model. Bar charts compare the adjusted R^2^ for predicting CDAI (left panel) and the area under the curve (AUC) for predicting complications (right panel) between the two models. The original model incorporates separate TG and HDL-C variables, while the ratio model uses the TG/HDL-C composite index. Error bars represent the 95% confidence intervals.

**Figure 6.**
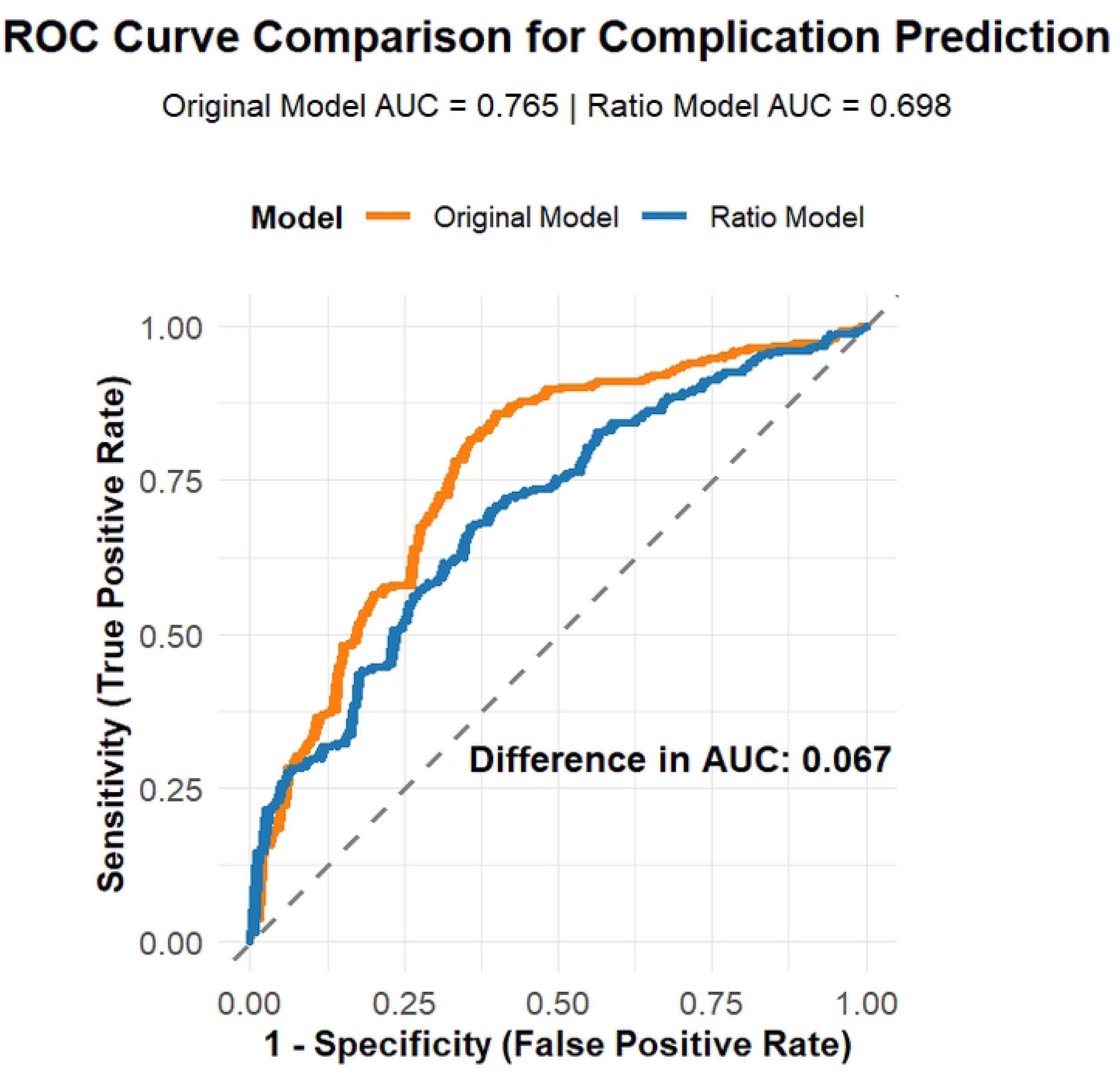
Comparison of receiver operating characteristic (ROC) curves for complication prediction. The ROC curve of the original model (AUC = 0.765) is directly compared with that of the TG/HDL-C ratio model (AUC = 0.698). The original model demonstrates superior discriminatory ability across most of the specificity range.

**Figure 7.**
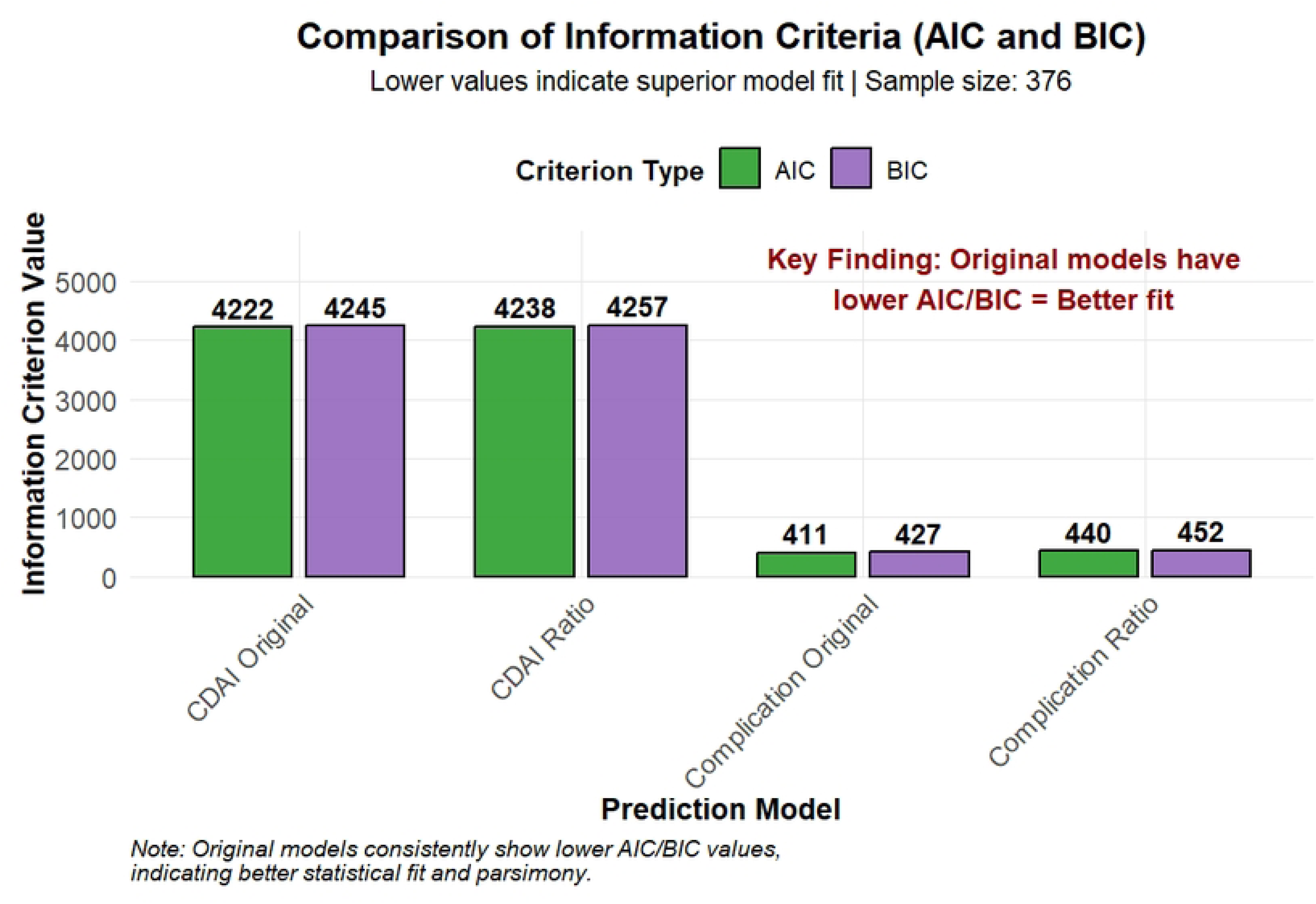
Comparison of model fit indices between the original and the TG/HDL-C ratio models. The Akaike Information Criterion (AIC) and Bayesian Information Criterion (BIC) values are shown for both the CDAI prediction (Panel A) and the complication prediction (Panel B) tasks. Lower AIC and BIC values indicate better model fit and parsimony, favoring the original model in both comparisons.

**Figure 8.**
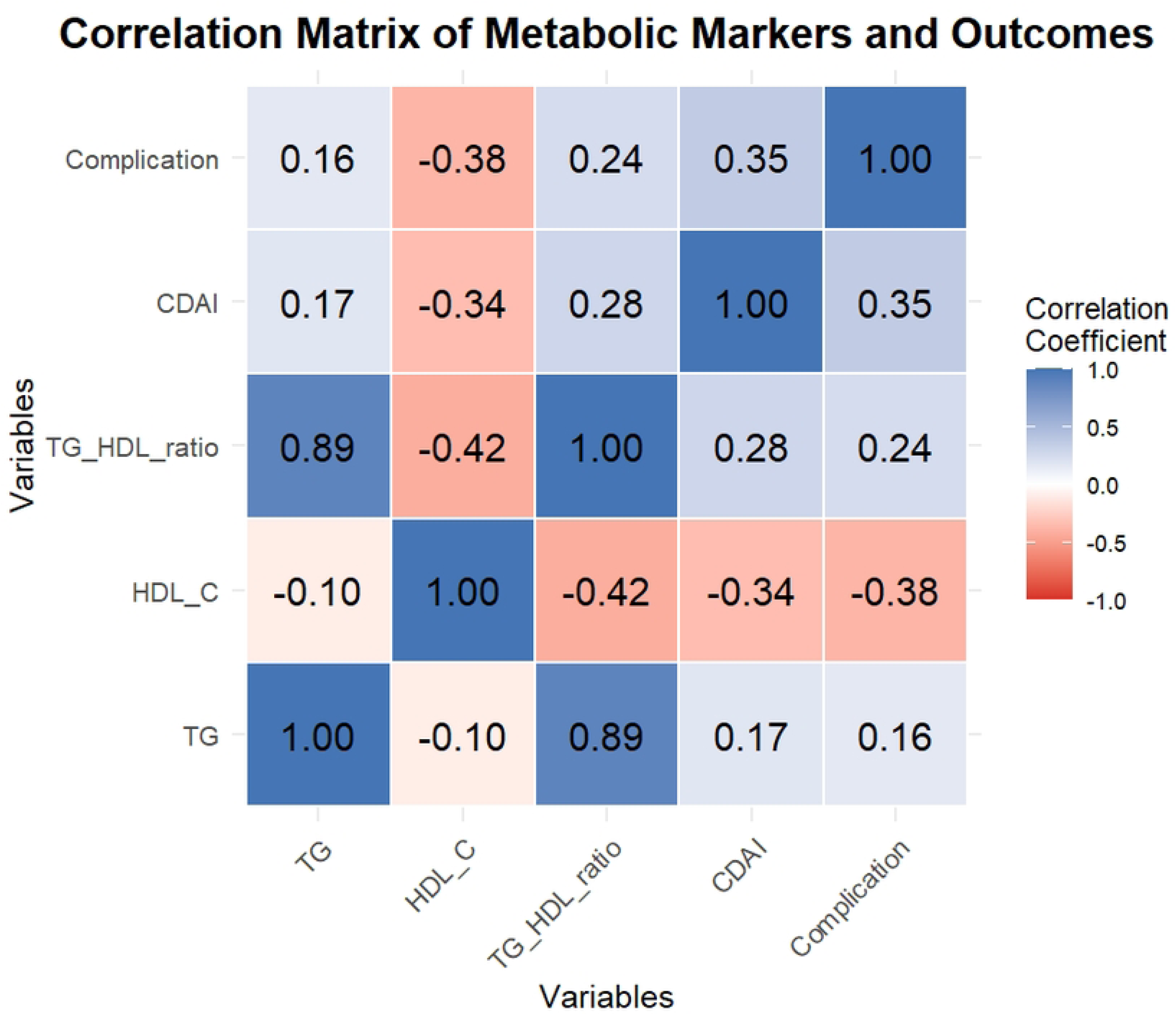
Correlation matrix heatmap of key metabolic markers and clinical outcomes. The heatmap visualizes Pearson correlation coefficients among Triglycerides (TG), high-density lipoprotein cholesterol (HDL-C), the TG/HDL-C ratio, Crohn’s Disease Activity Index (CDAI) score, and complication status. Red hues indicate positive correlations, blue hues indicate negative correlations, with color intensity proportional to the strength of the association.

## 3. Discussion

This study has delineated specific associations between metabolic disturbances and two pivotal clinical dimensions of CD. Regarding the clinical outcome reflected by CDAI scores, dyslipidemia—characterized by low HDL-C, low BMI, and a high triglyceride-to-HDL-C ratio—constituted a central driving factor, alongside the non-metabolic contributor of ESR. For the outcome of structural complication development, low HDL-C emerged as the strongest risk predictor, with undernutrition (low BMI) and lipotoxicity (high TG) also playing significant roles. The complication prediction model developed based on these insights demonstrated excellent discriminatory power (AUC=0.765) and calibration, providing a practical tool for the early identification of high-risk patients, with its robustness confirmed via bootstrap validation. As a common influencing factor, HDL-C plays an important role in both inflammatory activity and structural progression in CD.

We found that a Mendelian randomization study confirmed our conclusion that an elevated concentration of HDL-C with a genetic basis was significantly associated with a reduced risk of Crohn’s disease, with an odds ratio of 0.85^13^.High-density lipoprotein (HDL) plays a dual role in physiological systems. The significant anti-inflammatory effect of HDL is mainly mediated through two different pathways. It can directly bind to and neutralize endotoxins in the circulation, and can also alter the membrane composition of immune cells by extracting cholesterol from specialized lipid raft domains, thereby weakening the pro-inflammatory signal cascade. This latter action effectively suppresses signaling pathways associated with both Toll-like receptors (TLR) and T-cell receptor activation^14^. The HDL/S1P (sphingosine-1-phosphate) axis modulation of TGF-β might inhibit fibroblast activation, offering a potential mechanistic explanation for its influence on intestinal fibrogenesis and its predictive value for complication risk ^14^. HDL exhibits a variety of anti-inflammatory and antioxidant properties. An important part of this work further identified sphingosine-osteoporosis (S1P), a bioactive lipid medium carried on HDL, as the main molecular component responsible for these protective effects^15–17^. HDL3 subspecies synthesized by the intestinal epithelium (particularly in the ileum) are transported via the portal vein, bind to LPS-binding protein (LBP), and effectively sequester endotoxin, preventing TLR4 pathway activation in macrophages and thereby suppressing systemic inflammation ^18^. They also pointed out that the expression level of apolipoprotein A1 (apoA1) was significantly reduced in the ileum tissue of patients with Crohn’s disease, which was highly consistent with our findings that low HDL-C was associated with disease activity, suggesting that intestinal HDL synthesis disorder may be an important part of Crohn’s disease immune disorder^18^.

Recent studies indicate that elevating HDL-C not only alleviates intestinal inflammation via ATF3-dependent anti-inflammatory reprogramming of macrophages but also directly enhances intestinal epithelial barrier function ^19^. This indicates the protective potential of HDL-C in anti-inflammatory and structural damage. They also found that HDL-C levels inversely correlate with IBD inflammation and predict severity better than CRP ^19^. Relevant mechanisms have been previously reported; for instance, deletion of MCPIP1 in macrophages promotes M1 polarization and inhibits maturation into an anti-inflammatory phenotype via the ATF3-AP1S2 axis, exacerbating intestinal mucosal inflammation ^20^. Given that intestinal fibrosis involves fibroblasts and microbiota ^21^, and our finding links HDL-C to complications, a connection to fibrotic processes is plausible, warranting further investigation. Whether and how HDL directly regulates intestinal fibrosis by influencing fibroblasts or epithelial-mesenchymal transition awaits future investigation.

As a risk factor for complications, male gender may be related to the effect of sex hormones on the number, composition, and function of HDL ^8^, which explains why gender has lost its independent association after metabolic adjustment.

High TG, also an independent risk factor for both outcomes, highlights “lipotoxicity.” Research by a team from Peking University found that excess free fatty acids (e.g., palmitate) generated from hydrolysis in a high-TG environment can act as endogenous danger signals, directly activating the NLRP3 inflammasome in macrophages, leading to the release of key pro-inflammatory cytokines like IL-1β ^22^, causing systemic and even widespread inflammation. A high-fat diet pattern induces oxidative stress in the colon, enhances the permeability of the intestinal epithelial barrier, and promotes inflammatory responses^23^. Even short-term, acute high-fat intake impairs gut homeostasis, rapidly disrupting the intestinal barrier, altering gut microbiota, and significantly suppressing the secretion of IL-22—a key cytokine for barrier maintenance by group 3 innate lymphoid cells ^24^—leading to mucosal damage. The impairment of intestinal barrier integrity results in the direct contact between the mucosal surface and pro-inflammatory mediators. This exposure aggravates localized inflammation and concurrently disrupts the delicate microenvironment required for effective tissue regeneration, thereby establishing a foundation conducive to fibrotic complications. Separately, saturated fatty acids, including palmitate, inflict direct cytotoxic damage upon intestinal epithelial cells. Their mechanism involves the induction of endoplasmic reticulum stress coupled with compromised mitochondrial function, a sequence of events that culminates in programmed cell death and the consequent emergence of structural impairments within the physical barrier^25^. Autophagy alleviates ER stress^26^, but high-fat diets can impair barrier stress response via TLR4/NF-κ B-mediated autophagy ^27^. This reflects the inflammation-damage-repair failure cycle in high TG environments. Intestinally derived HDL protects the barrier ^28^, linking lipid disorders to barrier damage and suggesting TG-HDL interaction. Elevated TG often accompanies decreased HDL function, weakening the antioxidant and anti-inflammatory ability of intestinal microenvironment, promoting inflammatory cytokine infiltration and bacterial translocation. This imbalance in the metabolism-immunity-tissue repair interplay collectively drives the chronic inflammatory state in the gut, explaining why high TG associates with both short-term inflammatory activity (CDAI) and long-term penetrating/fibrotic complications (structural damage) in CD. Therefore, targeting lipid metabolism to reduce lipotoxicity may represent a potential novel strategy for improving CD prognosis.

TG/HDL-C ratio is a recognized indicator for identifying insulin resistance and atherogenic dyslipidemia ^29,30^. Our finding that separate TG and HDL-C models outperformed the ratio model suggests that in CD, TG and HDL-C may influence disease via relatively independent pathways. For instance, TG may lean more towards “lipotoxicity” and acute pro-inflammatory effects, while HDL-C may focus more on immunomodulation and barrier protection. The traditional composite ratio might obscure this complexity, whereas separate consideration captures more nuanced risk information, reflecting the complex specificity of CD.

We found BMI, as a common influencing factor for both outcomes, was closely linked to both CD inflammation and complication development. A study involving 601,009 participants concluded that obesity is associated with increased CD risk ^31^. However, there is a ‘higher BMI, better prognosis’ paradox in CD ^32^, which may be due to the inability of BMI to distinguish between fat and muscle mass (‘obesity paradox’). Research indicates that “sarcopenia” and “visceral obesity” are associated with poor prognosis in CD, while simple nutritional assessments are not^33^. BMI, as a general anthropometric index, has limitations and cannot assess visceral fat content; the traditional concept of “obesity” may mask sarcopenia risk ^34^. Recent clinical research suggests that in the specific context of CD, the simple label of “overweight/obesity” may not pose a major threat to core treatment outcomes ^35^. Low BMI may be due to long-term activity leading to impaired nutrition, because growth retardation precedes the diagnosis of CD in children^36^. Chronic undernutrition impairs immune and repair functions. The hazard may lie in visceral/ectopic fat (e.g., creeping fat), which promotes inflammation and fibrosis via local immunometabolic activities ^37^. In the context of obesity, adipocytes promote chronic inflammation by recruiting and stimulating CD8 + T cells and macrophages, a process partially mediated by β −2 microglobulin (B2M) 38. This mechanism’s insights remind us that the clinical management of Crohn’s disease patients should go beyond body mass index (BMI), including comprehensive nutritional assessment as well as evaluation of specific body composition indicators such as muscle mass and visceral adipose tissue.

Our research utilized the significant metabolic components identified through statistical analysis to develop a clinical predictive model for the occurrence of complications. Clinicians can use this tool to synthesize and graphically represent the possibility of complications based on metabolic profile data, providing stratified monitoring and personalized intervention plans. We also constructed a streamlined model for assessing disease activity, facilitating rapid evaluation of short-term inflammatory status through simple calculations. Therapeutic approaches aimed at increasing circulating levels or enhancing the functional capacity of high-density lipoprotein cholesterol should be prioritized in patients exhibiting low HDL-C. For individuals identified with elevated triglyceride-associated risk, clinical management must emphasize the control of lipotoxic effects to alleviate related inflammatory responses. When low body mass index is recognized as a contributing risk factor, implementing tailored nutritional interventions and structured rehabilitative care becomes critically important. These findings support the development of treatment for Crohn’s disease from a predominantly immunosuppression-centered model to a more comprehensive framework that integrates immune regulation, metabolic optimization, and nutritional support.

### 4.1 Limitations and Future Directions

Several study limitations need to be considered. Given the cross-sectional nature of this study, the relationships determined reflect those at a single point in time and no causal inferences can be drawn from them. Furthermore, the potential influence of residual confounding variables cannot be completely ruled out. Although the predictive model demonstrated robustness in internal validation, its universal applicability needs to be confirmed through external validation in independent cohorts or prospective longitudinal studies. To enhance the accuracy of predictions, future research can also examine the integration of more specific physiological indicators, such as directly evaluating HDL function or conducting detailed assessments of body composition. Prospective, multicenter cohorts would be valuable to validate and refine the presented models. Metabolomic profiling could also help identify disease-specific pathogenic lipid species and pathways in CD. For patients identified with high-risk metabolic phenotypes, interventional clinical trials could be designed to explore the adjunctive therapeutic potential of targeted metabolic modulation.

## 5. Conclusion

MetS components (low HDL-C, high TG, low BMI) are key determinants of clinical heterogeneity in CD. Dual-outcome analysis revealed specific association patterns, proposed novel mechanistic hypotheses (e.g., dual-hit of lipotoxicity, re-conceptualizing body composition), and developed practical clinical prediction models. Our work positions patient metabolic status centrally in CD management, providing new perspectives and tools for individualized risk assessment and future synergistic metabolic-targeted therapies.

## Acknowledgement

We thank all the patients who participated in this study. We are grateful to The First Affiliated Hospital of Guangxi Medical University for providing access to the clinical data.

## Data availability

The data underlying this article were provided by The First Affiliated Hospital of Guangxi Medical University under licence / by permission. Data will be shared on request to the corresponding author with permission of The First Affiliated Hospital of Guangxi Medical University.

## Author contributions

ICMJE criteria for authorship read and met: All authors. Agree with the manuscript’s results and conclusions: All authors. Designed the study: P.Y.X. Collected data and responsible for data integrity: H.S.S.&L.Y.H. Analyzed the data: L.Z.D. Wrote the first draft of the paper: P.Y.X. Revise it critically for important intellectual content:Q.S.Y.&J.H.X. Contributed to the writing of the paper:All authors. Interpretation of data, approved the final version of the manuscript: All authors. Guarantor of the article: P.Y.X.

